# Primary cilia and BBS4 are required for postnatal pituitary development

**DOI:** 10.1101/2025.07.15.664994

**Authors:** Kathryn M. Brewer, Katlyn K. Brewer, Nicholas C. Richardson, Jeremy F. Reiter, Nicolas F. Berbari, Mia J. Konjikusic

## Abstract

Primary cilia orchestrate several signaling pathways, and their disruption results in pleiotropic disorders called ciliopathies. Bardet Beidl syndrome (BBS), one such ciliopathy, provides insights into cilia function in many tissues. Using a mouse model of BBS, *Bbs4* knockout (*Bbs4^-/-^*), and conditional deletion of pituitary cilia, we found that adult *Bbs4^-/-^* and cilia-less pituitaries are hypoplastic and have increased gonadotroph populations. The developing *Bbs4*^-/-^ pituitary experiences mildly reduced Hedgehog signaling. Isolated *Bbs4*^-/-^ pituitary stem cells exhibited reduced Hedgehog signal responsiveness and expression of stem cell markers. These data demonstrate that cilia and BBS function are necessary for pituitary growth. We propose that altered pituitary cilia-mediated patterning contribute to physiological features of ciliopathies such as BBS.

**Summary statement:** A mouse model of BBS reveals that ciliary Hedgehog signaling is necessary for postnatal pituitary development.

## Introduction

The pituitary is considered the body’s master hormonal regulator, as it regulates many other organs (Alatzoglou et al., 2020). As an intermediary between the brain and peripheral organs, the pituitary responds to signals from the hypothalamus and releases hormones into the blood stream. Pituitary hormones help regulate, for example, the adrenals, mammary glands and gonads, to contribute to growth, reproduction, lactation and stress responses (Musumeci et al., 2015; Perez-Castro et al., 2012). Pituitary dysfunction can result in gigantism, acromegaly, dwarfism, delayed puberty, infertility, hypothyroidism, and metabolic issues (Owolabi et al., 2024).

The pituitary develops from Rathke’s pouch, an invagination of the oral ectoderm (Rizzoti, 2015). Subsequently, pituitary progenitors in Rathke’s Pouch give rise to multiple endocrine cell types (Bian et al., 2023; Davis et al., 2016; Kioussi et al., 1999; Treier et al., 2001). Morphogens (*e.g.*, Sonic Hedgehog (SHH)) and transcription factors (*e.g.*, Prop1) coordinate developmental patterning of this complex organ (Carreno et al., 2017; Davis et al., 2016; Sheng and Westphal, 1999; Treier et al., 2001, 1998; Treier and Rosenfeld, 1996).

All pituitary cell types are present at birth. However, postnatally, pituitary maintenance and expansion relies on a SOX2-expressing stem cell population within the marginal zone and anterior pituitary (Carbajo-Pérez and Watanabe, 1990; Pérez Millán et al., 2016; Taniguchi et al., 2001a, 2001b, 2000; Zhu et al., 2015). These SOX2 stem cells coordinate proliferation and expansion of the tissue. After the first few weeks of life, proliferation attenuates, but pituitary cellular composition changes in different physiological states (Rizzoti et al., 2013), such as puberty (Oishi et al., 1993) and pregnancy (Gonzalez et al., 1988). Despite evidence of cellular remodeling, little is known about the signaling pathways that regulate pituitary remodeling and homeostasis in postnatal life.

SHH is a morphogen that participates in the development of many different organs. During pituitary development, SHH is secreted from the oral ectoderm and is critical for the invagination of Rathke’s pouch and proliferation of Rathke’s pouch progenitors (Treier et al., 2001; Wang et al., 2010). SHH is also secreted in a subsequent step from the hypothalamus, where it specifies LHX3/4 pituitary progenitors (Carreno et al., 2017; Treier and Rosenfeld, 1996). Disruption of Rathke’s pouch development can lead to hypoplastic and mis-patterned pituitaries (Treier et al., 2001; Wang et al., 2010). In the adult pituitary, SHH is redeployed to regulate hormone production (Botermann et al., 2021; Pérez Millán et al., 2016). Additionally target genes of SHH signaling have been reportedly expressed downstream of SOX2+ stem cells in the pituitary (Pérez Millán et al., 2016; Sheridan et al., 2025). Disruption of this function can result in pituitary diseases like Pituitary Stalk interruption Syndrome or hypopituitarism (Aouchiche et al., 2025; Arnhold et al., 2015; Brauner et al., 2020).

HH signals in vertebrates are transduced through primary cilia, small, antenna-like microtubule-based projections on the surface of most cell types. The primary cilium transduces SHH signals by dynamically altering ciliary receptor composition (Corbit et al., 2005; Mukhopadhyay et al., 2013; Rohatgi et al., 2007). When SHH is present, it binds to its receptor, PTCH1, and causes its exit from the cilium and activation of GLI transcription factors which travel to the nucleus to induce SHH-responsive genes, such as *Ptch1* and *Gli1*. Ciliary defects can disrupt SHH signal transduction, altering the development of many organs including the pituitary (Lodge et al., 2023; Yoshida et al., 2024).

Several ciliopathies, including Bardet Biedl Syndrome (BBS), are characterized by developmental defects such as early pediatric obesity and cognitive impairment (Derderian et al., 2023; Mill et al., 2023; Reiter and Leroux, 2017). BBS is caused by mutations in proteins necessary for trafficking ciliary receptors into and out of the cilium in a signaling dependent manner (Brewer et al., 2023; Stubbs et al., 2023; Ye et al., 2018). BBS individuals experience disorders associated with altered pituitary function, including hypogonadism, hypothyroidism and hyperlactatemia (Taniguchi et al., 2001a, 2001b, 2000; Zhu et al., 2015). Some BBS individuals also exhibit hypoplastic pituitaries and hormonal imbalances (Guran et al., 2011). Moreover, knockout of any of the components of the BBSome complex, e.g. *Bbs4*, in mice causes altered body size, weight, bone length and gonadal size (Mykytyn et al., 2004), all of which can result from pituitary dysfunction (Caba et al., 2022). Given the connections between pituitary development, SHH signaling and ciliopathies, we investigated the function of cilia in pituitary formation, patterning and function.

## Results

### *Bbs4^-/-^* mice exhibit pituitary hypoplasia and increased gonadotroph proportions

To examine whether BBS4 is required for pituitary morphology, we histologically examined 8- week-old male and female *Bbs4^-/-^* and littermate control pituitaries (**Fig 1A** and **Supp.** Fig 1A, respectively). *Bbs4^-/-^* pituitaries were smaller (**Fig 1B**), consistent with our previous observations in *Bbs5^-/-^*pituitaries (Bentley-Ford et al., 2021). We did not observe any changes in the apportionment of the pituitary into its three lobes, the pars distalis (PD), pars intermedialis (PI), and pars nervosa (PN), in either males or females (**Fig 1B** and **Supp.** Fig. 1B). The cell density of the *Bbs4^-/-^*PD was reduced specifically in males (**Fig 1B**).

**Figure 1:**
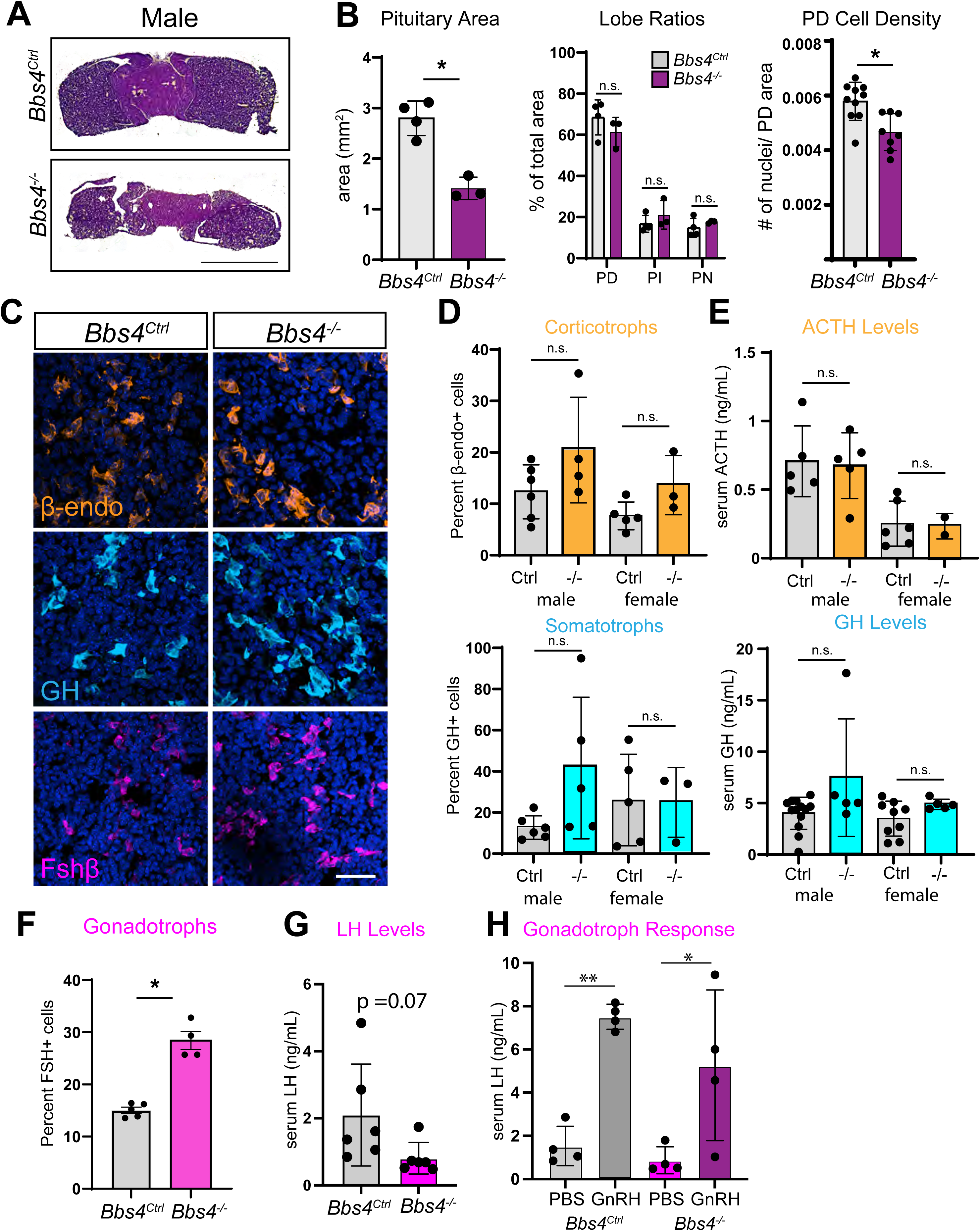
Pituitary hypoplasia and altered PD cell populations in adult *Bbs4* mutants. **A.** H&E staining of 8-week-old male pituitaries of mutants (*Bbs4^-/-^*) and controls (*Bbs4^Ctrl^*). Scale Bar 1 mm. **B.** Area and morphology analysis show *Bbs4^-/-^* mice have smaller pituitaries compared to *Bbs4^Ctrl^* control males (student’s t-test). There is no significant difference in the ratio of the 3 lobes of the pituitary (PD, PI, and PN) (one-way ANOVA)). The PD cell density is less in *Bbs4^-/-^*compared to *Bbs4^Ctrl^* control littermates (student’s t-test). **C.** Immunofluorescence staining of hormones in the PD for corticotropes (β-endo, orange), somatotrophs (GH, cyan), and gonadotrophs (FSH, magenta). Hoechst nuclei blue. Scale bar 50 µm. **D.** Quantification of the percent of corticotropes (β-endo) and somatotrophs (GH) show no significant differences between male and female Bbs4^-/-^ and Bbs4^Ctrl^ littermate controls (one-way ANOVA). **E.** Serum ACTH and GH of males and females show no significant differences (one-way ANOVA). **F.** Increase in the number of gonadotroph cells (FshB+) in *Bbs4^-/-^* compared to *Bbs4^Ctrl^* littermate controls (student’s t-test) **G.** Serum LH levels in male *Bbs4^-/-^* compared to *Bbs4^Ctrl^* controls (student’s t-test). **H.** Gonadotroph response to injection with GnRH in males measure via serum levels of LH which increased when compared to vehicle (PBS) injected animals (one-way ANOVA). ** indicates P < 0.01, * indicates P < 0.05, n.s. indicates not significant throughout. All animal n numbers indicated in graphs with a data point = 1 animal.

To determine if the reduction in mutant pituitary size and cell density was associated with changes in hormonal cell composition, we measured the proportion of different cell types in *Bbs4^-/-^* and control pituitaries using immunofluorescence (**Fig 1C**). The proportion of corticotrophs (ACTH, β- endo producing cells) and somatotrophs (GH, growth hormone producing cells) in the PD was unchanged (**Fig 1D**). In males, the percent of somatotrophs is highly variable in *Bbs4^-/-^* mutants compared to littermate controls whereas there are no significant changes or variability in female mutants (**Fig 1D**). To assess basal pituitary activity, we performed ELISA assays on serum from 8-week-old males and females to measure circulating hormones secreted by cells within the PD. Levels of adrenocorticotropic (ACTH) or growth hormone (GH) in *Bbs4^-/-^* mutants were unchanged (**Fig. 1E**).

Given that hypogonadism is a common, clinical feature of BBS and *Bbs4^-/-^*mice are subfertile (Mykytyn et al., 2004), we quantitated gonadotrophs (FSH, follicle stimulating hormone producing cells) within the PD (**Fig 1C**). Gonadotrophs were increased in *Bbs4^-/-^* males and females at both one month and 2 months of age (**Supp**. Fig 1C-D and **Fig 1F**). Basal serum levels of luteinizing hormone (LH) were unchanged in *Bbs4^-/-^*mice (**Fig. 1G**). To test if these mutant gonadotrophs were responsive to hypothalamic stimuli, we injected gonadotropin releasing hormone (GnRH) into adult male mice. GnRH stimulation similarly stimulated circulating LH in *Bbs4^-/-^* mice and in controls (**Fig 1H**). Together, these data indicate that, although *Bbs4^-/-^* pituitaries are hypoplastic, the corticotrophs, somatotrophs and gonadotrophs formed are functional.

### Cilia restrict gonadotroph specification

BBS4 is critical for cilia signaling through BBSome function but is dispensable for ciliogenesis (Mykytyn et al., 2004). To test whether cilia are important for later steps in pituitary development, we ablated pituitary cilia. More specifically, we deleted *Ift88*, encoding an intraflagellar transport component critical for ciliogenesis (Haycraft et al., 2007), using *Prop1-Cre* (Davis et al., 2016; Pérez Millán et al., 2016; Zhu et al., 2015). *Prop1-Cre* induced recombination of the <u>mTmG</u> reporter allele in the RP by E12.5 (**Supp.** Fig. 2A) (Muzumdar et al., 2007). *Prop1-Cre* also induced some recombination in the forebrain and olfactory epithelium, but not in every embryo, (**Supp.** Fig. 2B), as previously reported (Zhu et al., 2015).

*Prop1-Cre Ift88^fl/fl^* pituitaries exhibited no cilia (**Supp** **Fig. 2C**). Both male and female *Prop1-Cre Ift88^fl/fl^* were smaller in length and body weight than littermate controls by postnatal day (P28) (**Fig 2A-B**). *Prop1-Cre Ift88^fl/fl^* mice did not display altered P28 gonadal or pituitary size (**Fig 2C-D**, **Supp.** Fig 2D). *Prop1-Cre Ift88^fl/fl^* pituitaries exhibited normal concentrations of SOX2-expressing stem cells in the postnatal PD (**Fig 2E-G**). Strikingly, like *Bbs4^-/-^*pituitaries, *Prop1-Cre Ift88^fl/fl^* pituitaries exhibited increased density of FshB-expressing gonadotrophs already at one month of age (**Fig 2H-I**). Thus, primary cilia in the pituitary, like BBS4, are necessary to restrict specification of pituitary gonadotrophs post E12.5.

**Figure 2:**
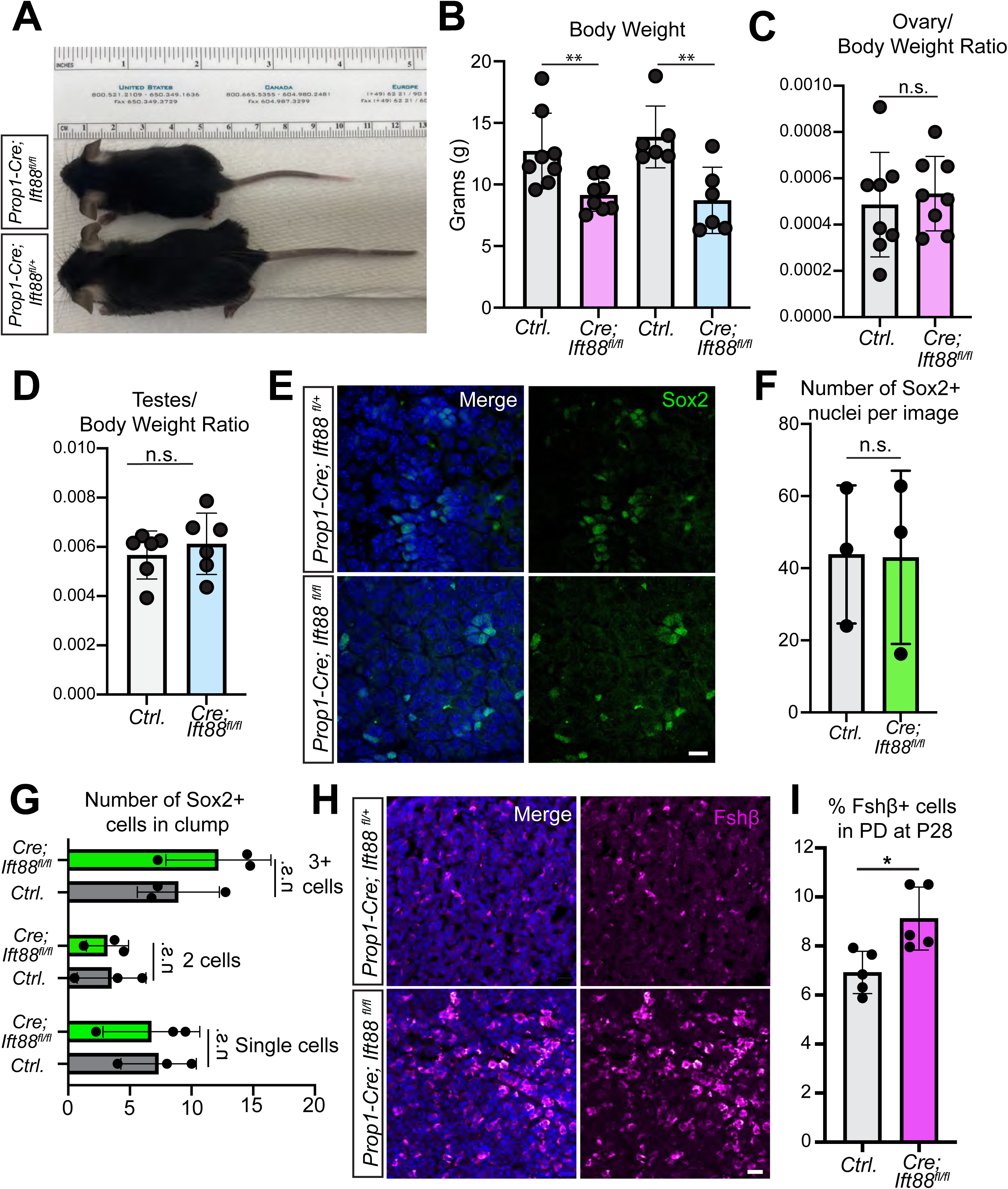
Conditional pituitary cilia ablation recapitulates *Bbs4* mutant pituitary phenotypes. **A.** Image of an adult conditional cilia mutant (*Prop1-Cre Ift88^fl/fl^*) and littermate control (*Prop1-Cre Ift88^fl/+^*). **B.** Body weights of *Prop1-Cre Ift88^fl/fl^* and littermate control (Ctrl) females (pink bars) and males (blue bars) at postnatal day 28 (P28). **C.** Ovary to body weight ratio and **D.** Testis to body weight ratio of mutants (*Prop1-Cr Ift88^fl/fl^*) compared to controls. **E.** Immunofluorescence of stem cells (SOX2, green) in *Prop1-Cre Ift88^fl/fl^* (*Cre;If88^fl/fl^*) and littermate controls (Ctrl). **F.** Quantification of SOX2+ cells in conditional mutants (*Prop1-Cre;Ift88^fl/fl^*) and littermate controls (*Ctrl*). **G.** Quantification of number of SOX2+ stem cells per rosette cluster in PD in mutants (*Cre;Ift88^fl/fl^*) and littermate controls (Ctrl.). **H.** Immunofluorescence for gonadotrophs (Fshβ, magenta) in P28 pituitaries in mutants (*Prop1-Cre;Ift88^fl/fl^*) and littermate controls (*Prop1-Cre;Ift88^fl/+^*). **I.** Quantification of the percentage of Fshβ+ cells in the PD of P28 mutant (*Cre;Ift88^fl/fl^*) and littermate controls (*Ctrl.).* All scale bars are 10µm and Hoechst nuclei blue. Student’s t-test ** indicates P < 0.01, * indicates P < 0.05, n.s. indicates not significant.

### Embryonic SHH signal transduction is mildly reduced in the absence of BBS4

We sought to determine if altered embryonic development accounts for the hypoplasia of *Bbs4^-/-^* pituitaries. As cilia orchestrate SHH signaling (Corbit et al., 2005; Mukhopadhyay et al., 2013; Rohatgi et al., 2007), and both SHH and primary cilia regulate early development of Rathke’s pouch (Lodge et al., 2023; Yoshida et al., 2024), we assessed ciliation and SHH signal transduction in embryos. We observed RP cilia at embryonic day (E) 10.5, a time point during which SHH signaling is active (Treier et al., 2001; Wang et al., 2010), by immunofluorescence for ARL13B, a ciliary marker, and CEP43, a marker of the basal body (**Fig 3A**) (Caspary et al., 2007). To understand if SHH signaling was reduced in *Bbs4^-/-^* mice, we performed fluorescent *in-situ* hybridization for the SHH target gene *Ptch1* and compared its expression to the control gene *Ubc*. *Ptch1* expression was mildly reduced in *Bbs4^-/-^* mice at E10.5 (**Fig 3B** and **C**).

**Figure 3:**
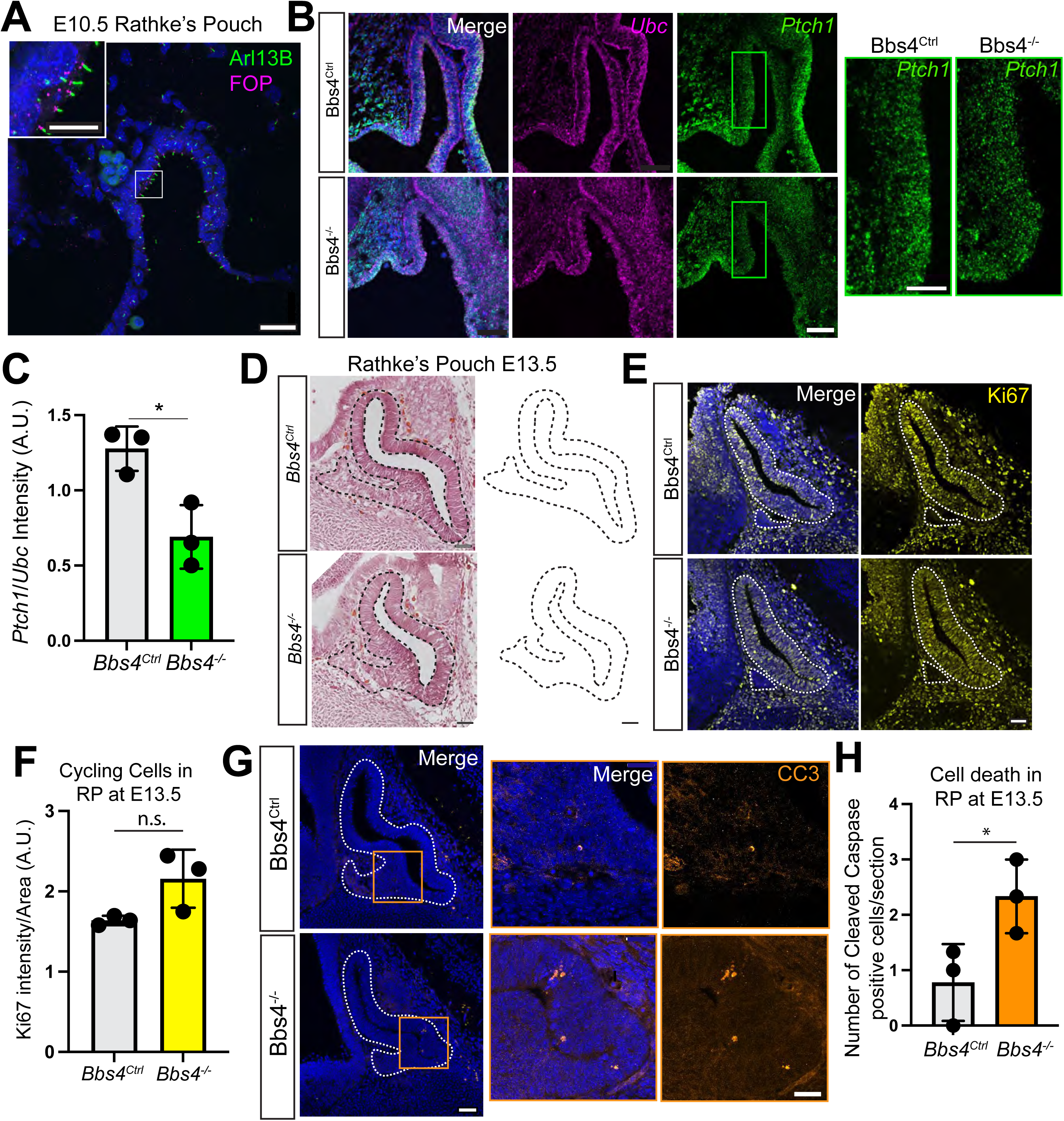
Hedgehog signal transduction is reduced in *Bbs4^-/-^* Rathke’s pouch. **A.** Immunofluorescence staining of cilia (ARL13B, green) and basal bodies (CEP43, magenta) in the RP at embryonic day 10.5 (E10.5). Scale bar 5 µm, Inset scale bar 2 µm. **B.** Fluorescence *in situ* hybridization of the RP at E10.5 in mutants (*Bbs4*^-/-^) and heterozygous (*Bbs4*^+/-^) littermate controls for SHH responsive (*Ptch1,* green) and internal control (*Ubc,* magenta) gene probes. Green box indicates inset for *Ptch1*. Scale bars 10 µm. **C.** Quantification of *Ptch1* probe fluorescence intensity compared to *Ubc* probe intensity in *Bbs4^+/-^* and *Bbs4*^-/-^ RP (P = 0.0170). **D.** H&E staining of staining of E13.5 RP in *Bbs4^-/-^*and *Bbs4^+/-^* controls. Scale bar 20 µm. **E.** Immunofluorescence staining of proliferative cells (Ki67, yellow) in *Bbs4^-/-^* and *Bbs4^+/-^* controls. Scale bar 50 µm. **F.** Quantification of Ki67 staining in *Bbs4^-/-^* and *Bbs4^+/-^* controls. **G.** Immunofluorescence staining for cell death using cleaved caspase three (CC3, Orange). Orange box indicates inset for CC3. Scale bars 50 µm and 10 µm for inset. **H.** Quantification of CC3 cells per animal in the RP at E13.5 in *Bbs4^-/-^* and *Bbs4^+/-^*controls. Dashed outlines indicate RP and Hoechst nuclei blue throughout. All animal n numbers indicated in graphs with a data point = 1 animal. (student’s t-test), * indicates P < 0.05, n.s. indicates not significant.

SHH signaling in early RP development drives cell proliferation and survival (Lodge et al., 2023; Treier et al., 2001; Wang et al., 2010; Yoshida et al., 2024). We performed H&E on E13.5 sections and observed no gross morphological changes (**Fig 3D**). To see if the slight reduction in SHH signaling drove any defects in cell cycle or cell survival, we stained E13.5 sections of Rathke’s Pouch for Ki67 for overall cell cycle states and Cleaved Caspase 3 (CC3) for cell death. The proportion of Ki67-expressing cells was similar in E13.5 *Bbs4^-/-^* and control pituitaries (**Fig 3E-F**). There was a mild increase in CC3-positive cells in *Bbs4^-/-^* pituitaries compared to controls (**Fig 3G-H**).

### Late gestational pituitary differentiation does not require BBS4

SHH signaling during RP development contributes to generating the different hormone-producing cell types (Carreno et al., 2017; Lodge et al., 2023; Treier et al., 2001; Wang et al., 2010; Yoshida et al., 2024). A dorsal/ventral gradient of morphogens patterns the pituitary into the *Prop1*- expressing anterior pituitary primordium, *Islet1*-expressing ventral domain, and *Pax6-*expressing dorsal domain (Bian et al., 2023) (**Fig 4A**). To determine if pituitary patterning is disrupted by earlier decreases in SHH signaling during RP development, we assessed expression of *Islet1*, *Prop1* or *Pax6* at E13.5 (**Fig 4B**-**C**). Deletion of BBS4 did not dramatically alter expression. Moreover, assessing cell types in pituitaries at the end of embryonic development, E17.5, revealed no differences in the number of SOX2-expressing stem cells, LH-expressing gonadotrophs, or ACTH-expressing corticotrophs in *Bbs4^-/-^* pituitaries (**Fig 4D-K**). Given that adult *Bbs4* mice do display disrupted cell types (**Fig 1**), we conclude that the mild decrease in SHH signal transduction during embryonic development does not disrupt overall pituitary patterning or the induction of corticotrophs or gonadotrophs.

**Figure 4:**
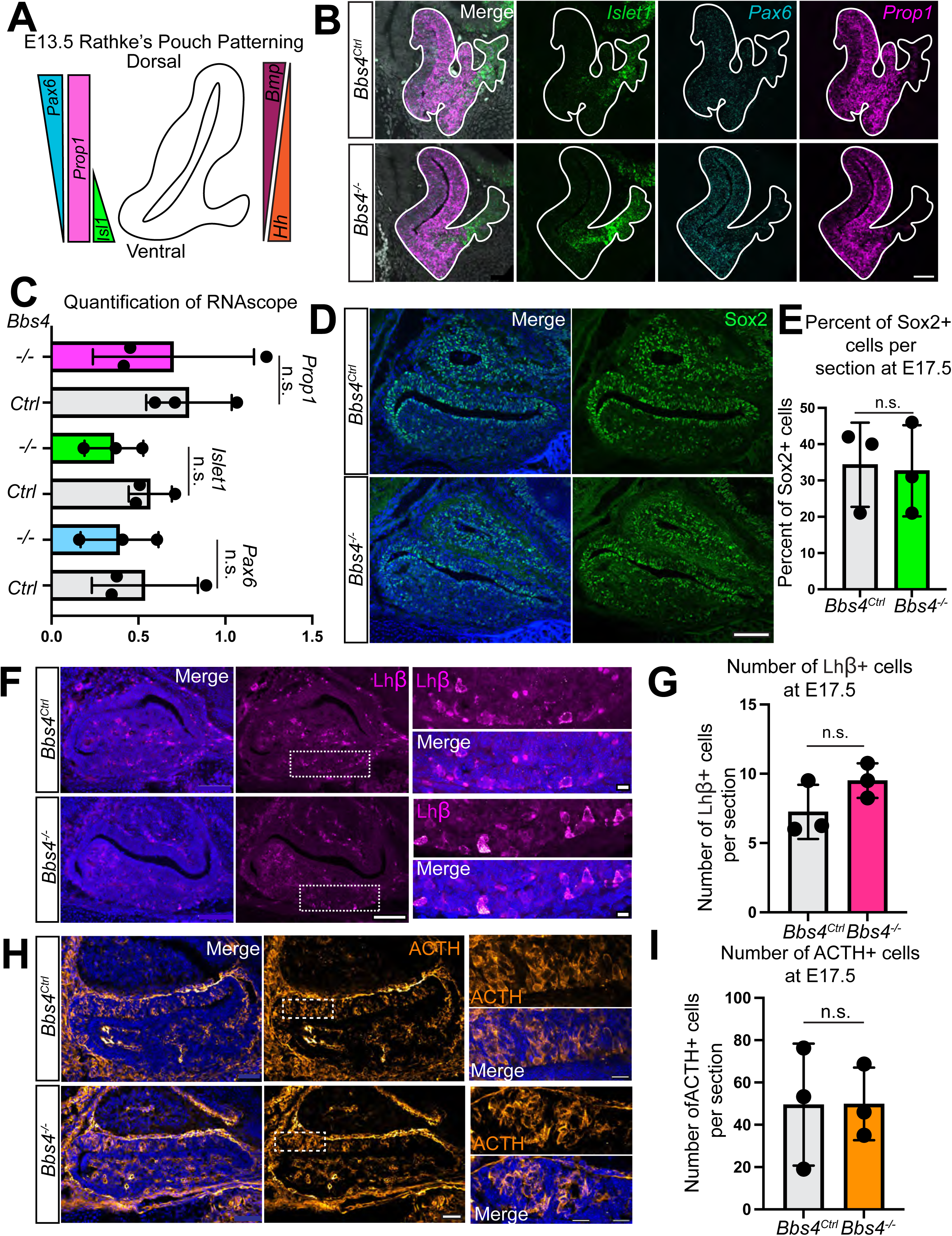
Cell differentiation is unaffected in *Bbs4^-/-^* pituitary. **A.** Schematic of morphogen gradients (Bmp, SHH), patterning, and cell fates markers in the dorsal ventral plane of the E13.5 RP (Prop1, Isl1, Pax6). **B.** Fluorescence *in situ* hybridization for key patterning genes merged and individually (Hoescht: grey, *Islet1:* green, *Pax6*: cyan, and *Prop1,* magenta) Scale bar 50 µm. **C.** Quantification of in situ hybridization probe fluorescence intensity comparing *Bbs4^-/-^* and *Bbs4^+/-^*control embryos. **D.** Immunofluorescence staining for pituitary stem cells (SOX2, green) in the RP of E17.5 embryos of *Bbs4^-/-^* and *Bbs4^+/-^* controls. Scale bar 200 µm. Hoechst nuclei blue. **E.** Quantification of average number of SOX2+ cells in *Bbs4^-/-^* and *Bbs4^+/-^* per embryo. **F.** Immunofluorescence staining for early gonadotrophs (LH, magenta) in the RP of E17.5 embryos of *Bbs4^-/-^* and Bbs4^+/-^ controls. Hoechst nuclei blue. Scale bar 200 µm and 10 µm for insets. **G.** Quantification of LH+ cells in *Bbs4^-/-^* and *Bbs4^+/-^* controls. **H.** Immunofluorescence staining for early corticotropes (ACTH, orange) in the RP of E17.5 embryos of *Bbs4^-/-^* and *Bbs4^+/-^* controls. Hoechst nuclei blue. Scale bar 200 µm and 10 µm for insets. **I.** Quantification of ACTH+ cells in *Bbs4^-/-^* and Bbs4^+/-^ controls. Dashed outlines indicate RP. All animal n numbers indicated in graphs with a data point = 1 animal. Student’s t-test, * indicates P < 0.05, n.s. indicates not significant.

### Ciliary signaling is necessary for postnatal SOX2-expressing stem cell maintenance and hedgehog responsiveness

During the first weeks of life after birth, pituitary stem cells continue to grow the pituitary gland (Pérez Millán et al., 2016; Taniguchi et al., 2001a, 2001b, 2000; Zhu et al., 2015). Later in adulthood, pituitary cellular composition is thought to change to meet different physiological demands(Gonzalez et al., 1988; Oishi et al., 1993). We hypothesized that BBS4 functions postnatally to control pituitary growth and composition. To test this hypothesis, we assessed the ciliation of control and *Bbs4^-/-^* adult PD (**Fig 5A**). Loss of BBS4 did not alter ciliation or cilia length in either male or female mice (**Fig 5B**).

**Figure 5:**
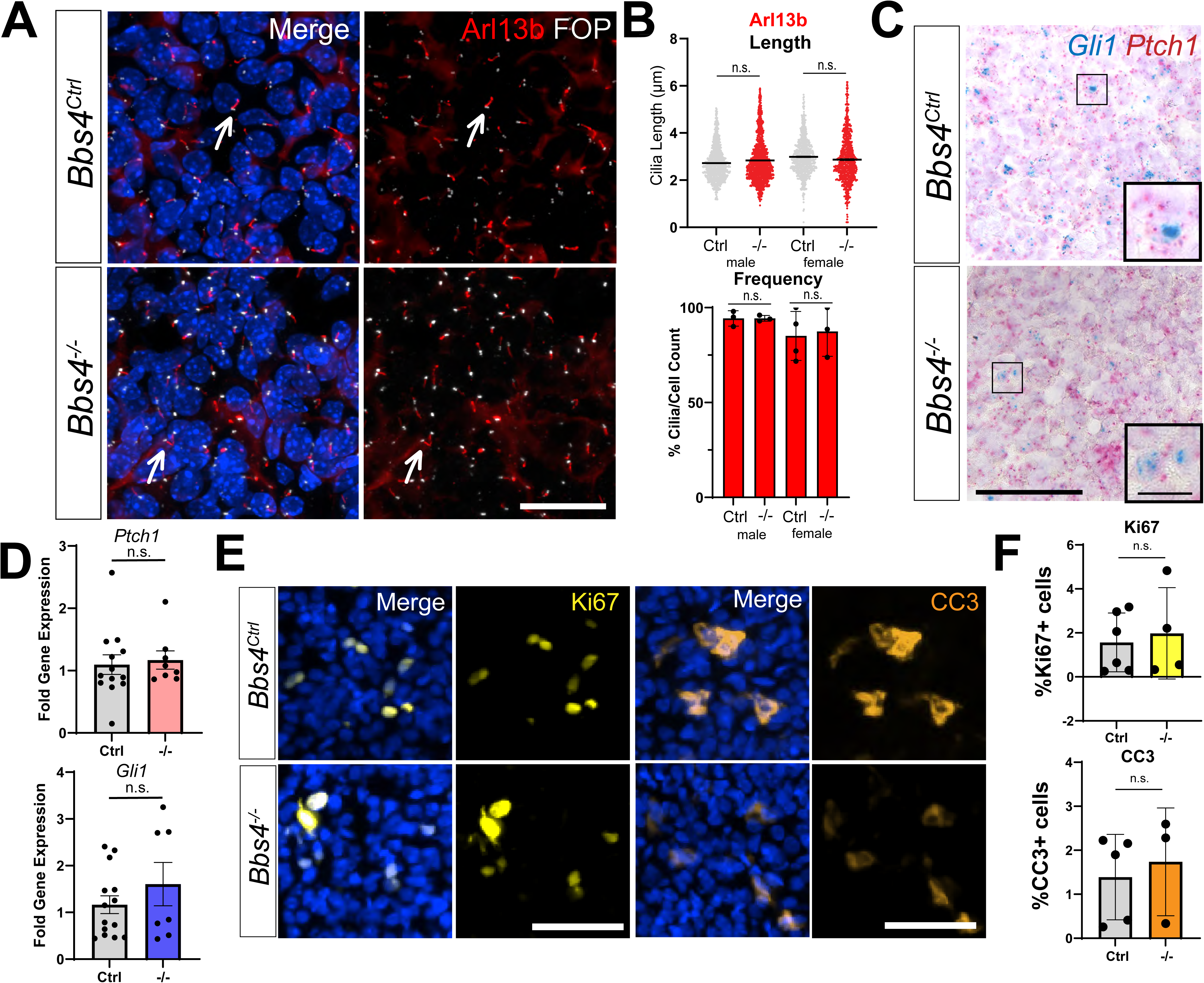
Assessment of adult *Bbs4* mutant pituitary PD. **A.** Immunofluorescence staining of cilia (ARL13B red) and basal bodies (FOP, white) in the PD of adult animals. Scale bar, 50 µm. **B.** Cilia analysis of Arl13b cilia length and frequency (One-way ANOVA). **C.** Colorimetric *in situ* hybridization in the PD of adult mutant (*Bbs4^-/-^*) and control (*Bbs4^Ctrl^*) littermates for SHH responsive probes (*Gli1*, blue and *Ptch1,* red). Hematoxylin counterstain and scale bars 50 µm. **D**. Graphs of RT-qPCR analysis of adult whole pituitary RNA for *Gli1* and *Ptch* in controls (*Ctrl*) and mutants (*-/-*). (student’s t-test). **E.** Ki67 (cyan) and CC3 (yellow) staining in the PD of *Bbs4* control and mutant animals. Hoechst nuclei blue. Scale bars 50 µm. **F.** Analysis of Ki67 and CC3 positive cells in the PD *Bbs4* control and mutant animals. Student’s t-test, ** indicates P < 0.01, * indicates P < 0.05, n.s. indicates not significant throughout. All animal n numbers indicated in graphs with a data point = 1 animal.

To assess whether BBS4 contributes to SHH signal transduction in the adult, we assessed *Ptch1* or *Gli1* expression in the PD of P56 male and female animals by *in situ* hybridization (**Fig 5C**) and RT-qPCR. We did not detect BBS4-dependent differences by either method (**Fig 5D**). Additionally, adult *Bbs4^-/-^* PDs exhibited no change in the number of proliferating or apoptotic cells (**Fig 5E-F**).

Although there were no broad changes to pituitary SHH signaling in *Bbs4* mutants, we hypothesized that specific subpopulation of pituitary cells may rely on BBS4 for SHH signal transduction. To test this hypothesis, we stained P56 pituitaries for the stem cell marker SOX2 (**Fig 6A**) and assessed the expression of pituitary *Sox2* using RT-qPCR. Neither the proportion of SOX2-expressing cells nor the expression of *Sox2* nor the clustering of SOX2-expressing cells was altered in *Bbs4^-/-^*pituitaries (**Fig 6B,C**).

**Figure 6:**
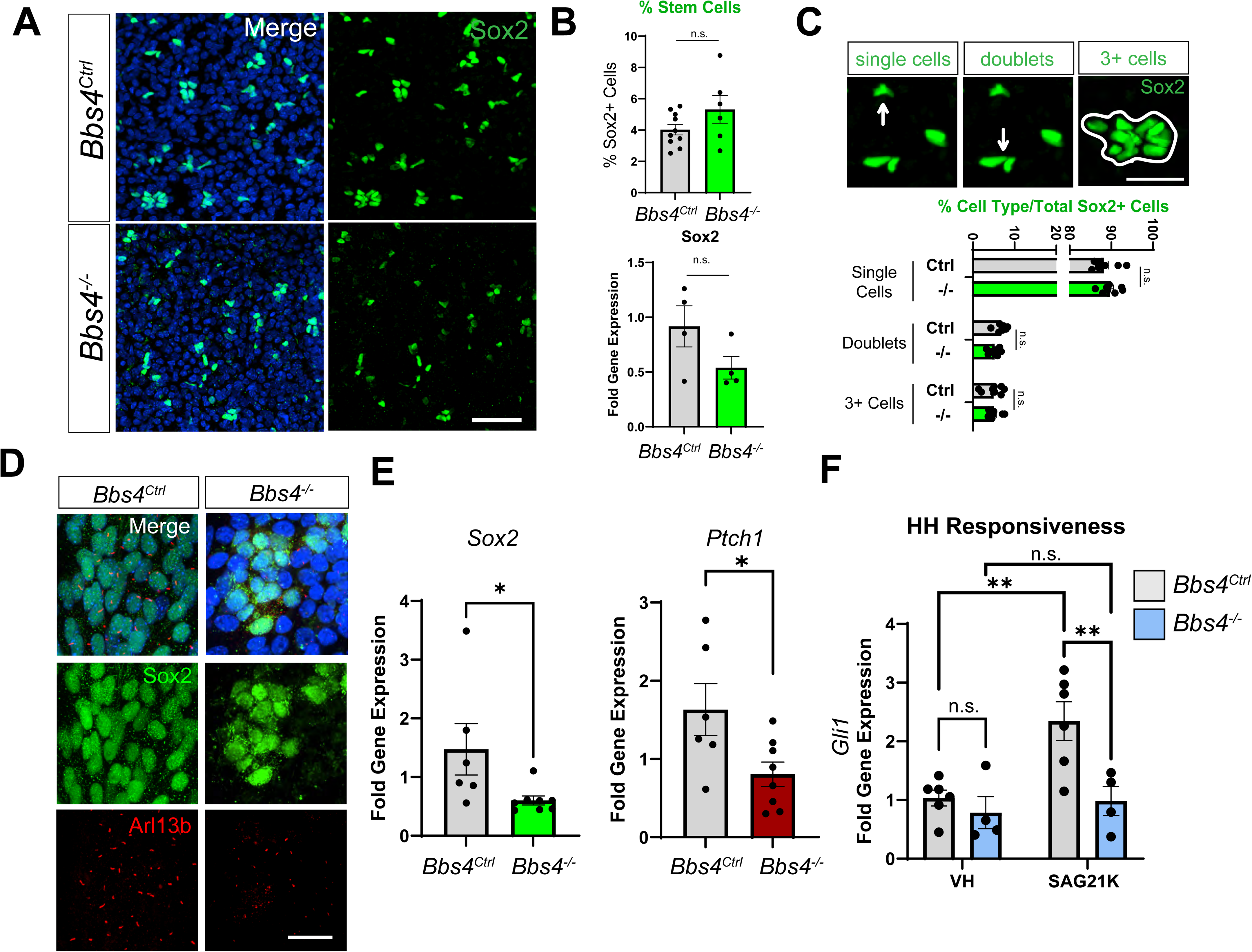
*Bbs4* mutant pituitary stem cells exhibit reduced Hedgehog signal transduction. **A.** Immunofluorescence staining for pituitary stem cells (SOX2, green) in the adult PD of *Bbs4^-/-^* mutants and *Bbs4^Ctrl^* controls. Hoechst nuclei blue. Scale bars 50 µm. B. Analysis of the number of SOX2+ cells and RT-qPCR expression of SOX2 in whole pituitary samples (student’s t-test). **C.** Immunofluorescence images and analysis of cluster sizes: single cells, doublets, and 3+ cells. (one-way ANOVA). Scale bar 10 µm. **D.** Immunofluorescence in mutant (*Bbs4^-/-^*) and control (*Bbs4^Ctrl^*) cultured pituitary stem cells (SOX2+, green) are ciliated (ARL13B, red). Scale bar *25* µm. **E.** RT-qPCR analysis of 7 DIV cultures for *SOX2* and *Ptch1* expression level in *Bbs4^-/-^* and *Bbs4^Ctrl^* cells (*student’s t-test*). **F.** Assessment of SHH responsiveness of *Bbs4^-/-^* and *Bbs4^Ctrl^* cells by RT-qPCR quantification of *Gli1* expression upon SHH activation with SAG21K compared to vehicle (VH) in 3 DIV stem cell cultures (*one-way* ANOVA). ** indicates P < 0.01, * indicates P < 0.05, n.s. indicates not significant throughout. All animal n numbers indicated in graphs with a data point = 1 animal. Hoechst nuclei blue throughout.

To further assess the potential altered signaling capacity of stem cells in the postnatal PD, we isolated and cultured stem cells from adult control and *Bbs4^-/-^* pituitaries (**Fig 6D**). Cultured *Bbs4^-/-^* stem cells exhibited decreased *Sox2* and *Ptch1* expression (**Fig 6E**). Stimulating with a SHH pathway activator, SAG21K, revealed that *Bbs4^-/-^* stem cells showed an attenuated ability to induce *Gli1* (**Fig 6F**). These results indicate that *Bbs4^-/-^*pituitary stem cells have a reduced capacity to respond to hedgehog signals.

## Discussion

We found that BBS4 and primary cilia are important for pituitary growth and the restriction of gonadotroph formation postnatally. In the absence of BBS4 or pituitary primary cilia, pituitaries are small and show increased proportions of gonadotrophs. Gonadotrophs in *Bbs4* mutants remain responsive to hormonal stimuli. We propose that although BBS is linked to hypogonadism, it is unlikely that its origin is defective pituitary gonadotroph function.

Primary cilia and SHH signaling have well-established roles in the early embryonic development of Rathke’s pouch, particularly in driving progenitor proliferation and organizing spatial patterning (Carreno et al., 2017; Lodge et al., 2023; Treier et al., 2001; Wang et al., 2010; Yoshida et al., 2024). Embryonic SHH signal transduction in Rathke’s pouch was mildly reduced in *Bbs4^-/-^* pituitaries and accompanied by an increase in cell death at E13.5. We propose that increased cell death contributes to *Bbs4* mutant pituitary hypoplasia. Mildly altered embryonic SHH signal transduction in *Bbs4* mutants was not accompanied by altered patterning in perinatal pituitaries, leading us to conclude that pituitary disruption in these mutants is likely to be caused by post- natal changes.

To further probe the temporal requirement for primary cilia function in pituitary development, we ablated cilia at E11.5-13.5 using *Prop1-Cre*. *Prop1-Cre* drives recombination in all pituitary hormone progenitors at E11.5-13.5 (**Supp.** Fig 2B, Davis et al., 2016; Zhu et al., 2015). *Prop1- Cre;Ift88^fl/fl^* mice were small with increased FshB-expressing gonadotroph proportions in their pituitaries at P28. Since SHH signaling is active in the postnatal pituitary (**Fig 5-6**), and HH signal transduction components are expressed by the SOX2-expressing progenitors in the pituitary (Pérez Millán et al., 2016; Sheridan et al., 2025), we propose that SHH signal transduction through pituitary primary cilia is necessary for pituitary growth and limiting the number of gonadotrophs.

SOX2-expressing stem cells undergo dramatic reshaping and expansion during the early postnatal period, contributing to the structural and functional maturation of the pituitary gland (Pérez Millán et al., 2016; Taniguchi et al., 2001a, 2001b, 2000). Postnatal stem cell populations specifically drive gonadotroph expansion between P7 and puberty (Sheridan et al., 2025). In considering the origins of the expanded gonadotroph populations in *Bbs4* mutants, we propose that the reduced ability to transduce SHH signals may contribute to overproduction of the gonadotroph fate. It is plausible that primary cilia and *Bbs4*-dependent ciliary signaling is required for proper maintenance of the pituitary proliferative population (Botermann et al., 2021; Sheridan et al., 2025).

Bardet-Biedl syndrome (BBS) is a ciliopathy wherein ciliary signaling is partially disrupted. We found that cilia and SHH signaling play important roles in postnatal pituitary homeostasis, potentially through regulation of SOX2-expressing stem cells. Our findings suggest that the BBSome is not critical for embryonic patterning or cell differentiation but the BBSome through primary cilia is critical for postnatal pituitary growth and restraint of gonadotroph formation.

## Materials and methods

### Mice

All procedures were approved by the Institutional Animal Care and Use Committee at Indiana University-Indianapolis and the University of California, San Francisco. Adult Bbs4 mice were obtained from The Jackson Laboratory (B6.129-*Bbs4^tm1Vcs^*/J Strain # 010728) (Mykytyn et al., 2004). *Prop1-Cre* (Zhu et al., 2015) mice were obtained from Xiaoyan Zhu at the Salk Institute. The *Ift88^fl^*allele was obtained from JAX (B6.129P2-*Ift88^tm1Bky^*/J, Strain #022409)(Haycraft et al., 2007). Unless specified, all experiments were conducted in male and female control and mutant animals. *Bbs4* mice at UCSF were maintained on a CD1 Charles River background. All other strains were maintained on a C57B6J/129 mixed background. Mice were housed on a standard 12 h light/dark cycle with *ad libitum* food and water.

### GnRH injections

3-month-old *Bbs4* male mice were IP injected at 150ng/kg with GnRH (Bachem Americas Inc., Cat #4033013) or PBS vehicle as described(Clarkson et al., 2017). Terminal blood collections for serum analysis were performed 15 minutes post injection. Whole blood was allowed to coagulate at room temperature for 90 minutes, then spun down at 2000 x G for 15 minutes and stored at - 20°C. ELISAs were performed by University of Virgina ligand Core for LH ELISAs.

### Fixation and tissue processing

Mice were anesthetized with a 0.1 ml/10 g body weight dose of 2.0% tribromoethanol (Sigma-Aldrich) and transcardially perfused with PBS, followed by 4% paraformaldehyde in PBS (catalog #15710, Electron Microscopy Sciences). Pituitaries were then isolated and postfixed in 4% paraformaldehyde for 30 minutes at 4°C and then cryoprotected using 30% sucrose in PBS for 16–24 hours. Pituitaries were embedded in optimal cutting temperature compound (catalog #4585, Thermo Fisher Scientific) and cryosectioned at 15 µm for analysis.

### Hematoxylin and Eosin staining

H&E staining on 15µm for adult and embryonic pituitary tissue sections was performed by Indiana University School of Medicine Histology Core.

### Immunofluorescence

Sections were washed with PBS for 5 minutes twice, then permeabilized and blocked in a PBS solution containing 1% BSA, 0.3% Triton X-100, 2% (v/v) donkey serum, and 0.02% sodium azide for 30 minutes at room temperature. For Ki67 staining, an addition wash in Antigen Unmasking Solution (H-3300, Vector Laboratories, Inc.) for 1 hour at 37°C was performed prior to the 30- minute blocking step. Sections were then incubated with primary antibodies (see Table 1.) in blocking solution overnight at 4°C. Sections were then washed with PBS (5 minutes wash twice) and blocking solution (5 minutes wash three times) before incubating with secondary antibodies for 1 hour at room temperature. All primary and secondary solutions were made in the blocking solution described above. Slides were then washed in PBS and stained with Hoechst nuclear stain (catalog #H3570, Thermo Fisher Scientific) for 5 minutes at room temperature. Coverslips were mounted using SlowFade Diamond Antifade Mountant (catalog #S36972, Thermo Fisher Scientific) and sealed with nail polish before imaging.

|  | Antibody | Catalog No. | Dilution |
| --- | --- | --- | --- |
| Cilia Analysis | ADCY3 | CPCA-ACIII | 1:1000 |
|  | ARL13B | ABIN1304543 | 1:300 |
|  | FOP | 11343-1-AP-20UL | 1:500 |
| Cell Analysis | GH | sc-374266 | 1:500 |
|  | LH/FSH | Gift from Buffy Elsworth, NIDDK antibodies | 1:300 |
|  | β-endorphin | H-022-33, Phoenix Pharmaceuticals | 1:1000 |
|  | Sox2 | Ab97959, Abcam | 1:1000 |
|  | Ki67 | Ab16667, Abcam | 1:1000 |
|  | Cleaved Caspase 3 | Antibody #9661, Cell Signaling Technology | 1:500 |

### *In situ* hybridization

Tissue cryosections (15 μm) were collected using our fixation and preparation protocol above and then prepped and pretreated with 4% PFA for 16 hours at 4°C, followed by part 2 of protocol TN 320534 (Advanced Cell Diagnostics (ACD), Newark, CA, USA) as described. Following tissue preparation, the detection of transcripts was performed using an RNAscope 2.5 HD Duplex Detection Kit (Chromogenic) User Manual Part 2 (ACDBio Document 322500-USM, Newark, CA, USA). The slides were assayed using a probe specific to *Gli1*, *Prop1*, *Islet1*, *Pax6*, and *Ptch1* transcripts (ACD), and counterstained with hematoxylin, dehydrated, and mounted using VectaMount (Vectorlabs, Burlingame, CA, USA), or counterstained with Hoechst and mounted with Prolong diamond for confocal fluorescence imaging. Slides with a positive control probe (PPIBC1/POLR2A; catalog no. 321651) and negative control probe (DapB; catalog no. 320751) were run with each experiment. At least 3 animals were analyzed for each group.

### RT-qPCR

Total RNA from whole pituitaries was isolated, cDNA was prepared, and quantitative real-time PCR was performed as described previously (Bansal et al., 2019). Assays-on-Demand Gene Expression Probes (Applied Biosystems) were as follows: *Ptch1* Mm00436026_m1; *Gli1* Mm00494654_m1; *Sox2* Mm03053810_s1. Ct values were normalized to *β-actin* Mm02619580 for whole pituitary collections and *Tbp* Mm01277042_m1 for cultured cells, relative expression was calculated by the ΔΔCt method.

### ELISAs

Blood was collected from 8-week old and 12-week old control and mutant mice in an EDTA tube and serum was isolated from the sample by being spun down at 10000 x G for 15 minutes and stored at –80 °C. Serum was then used to measure circulating growth hormone (GH), adrenocorticotropic hormone (ACTH), luteinizing hormone (LH) and follicle-stimulating hormone (FSH) using the following assay kits: GH ELISA (ABIN6956350) and ACTH ELISA (ab263880). Samples were ran through a SpectraMaxM5 plate reader from Molecular Devices immediately after assay completion. LH and FSH ELISAs were performed by the UVA ligand core.

### In vitro Stem Cell Isolation and Analysis

Sox2 positive stem cells were isolated from adult Bbs4 control and mutant animals. Briefly, the PI and PN were removed from pituitaries isolated under a dissecting scope. The remaining PD was then cultured, and stem cells were selected for growth through culturing conditions as previously described(Shintani and Higuchi, 2022). Stem cells were left to grow for 7 days with media being changed every 3 days. On day 7, cells were either fixed and assessed for stem cell properties and cilia, taken for RT-qPCR,or differentiated under conditions as described(Shintani and Higuchi, 2022). On day 3 in culture, stem cells were treated with either vehicle (EtOH) or 10nM SAG21K (Cat. No. 5282, Tocris BioTechne) for 3 hours. Treatments were then removed, and RNA was isolated for RT-qPCR. -qPCR.

### Imaging

All immunofluorescence and *in situ* images for analysis were acquired using a Nikon Ax confocal microscope or a Zeiss LSM 800 confocal microscope using water 20X (cell types) or air 20x (Cell types) and 40X (cilia analysis and E13.5 RNAscope analysis), or 63x for some cell type and E10.5 section analysis. Representative images of the entire pituitary were captured and stitched using a water 20X. H&E were captured using Leica ICC50 Brightfield scope.

### Image and Statistical Analysis

Computer-assisted cilia and cell analysis were performed using the Nikon artificial intelligence 5.30.06 software module as previously described(Bansal et al., 2021; Brewer et al., 2024, 2023). Threshold.ai was used to identify cilia and hormone positive cells as well as Hoechst positive cells. The area of the PD in each image was also measured for appropriate data analyses. Fiji (i.e. ImageJ) was used to assess and analyze H&E images for pituitary size and lobe ratio measurements. GraphPad Prism was used for statistical analysis with all tests and values reported in legends.

## Acknowledgements

We thank Buffy Elsworth, PhD, for the kind gift of the LH and FSH antibodies from the NIDDK Hormone Peptide Program. We also thank Michael G. Rosenfeld from UCSD and Xiaoyan Zhu from the Salk Institute for the kind gift of the *Prop1-Cre* mice. We thank the University of Virginia Ligand Core for performing the ELISAs on serum LH. We also thank the Reiter labs and the Berbari labs for thoughtful discussion and input.

## Competing interests

JFR is a cofounder of a BridgeBio-funded company and Renasant Bio.

## Funding

This work was supported by F31DK142351 (KMB), IU Indianapolis University Graduate Fellowship (KMB and KKB), the Beckman Scholars Program (NCR), R01DK114008 (NFB and JFR) and R01AR054396 and R01HD089918 (JFR), T32DK007418 (MJK) and A.P Giannini Postdoctoral and Leadership Award (MJK).

## Data and resource availability

Data and resource availability: All relevant data and resource can be found within the article and its supplementary information.

---

**Supplemental Figure 1.**
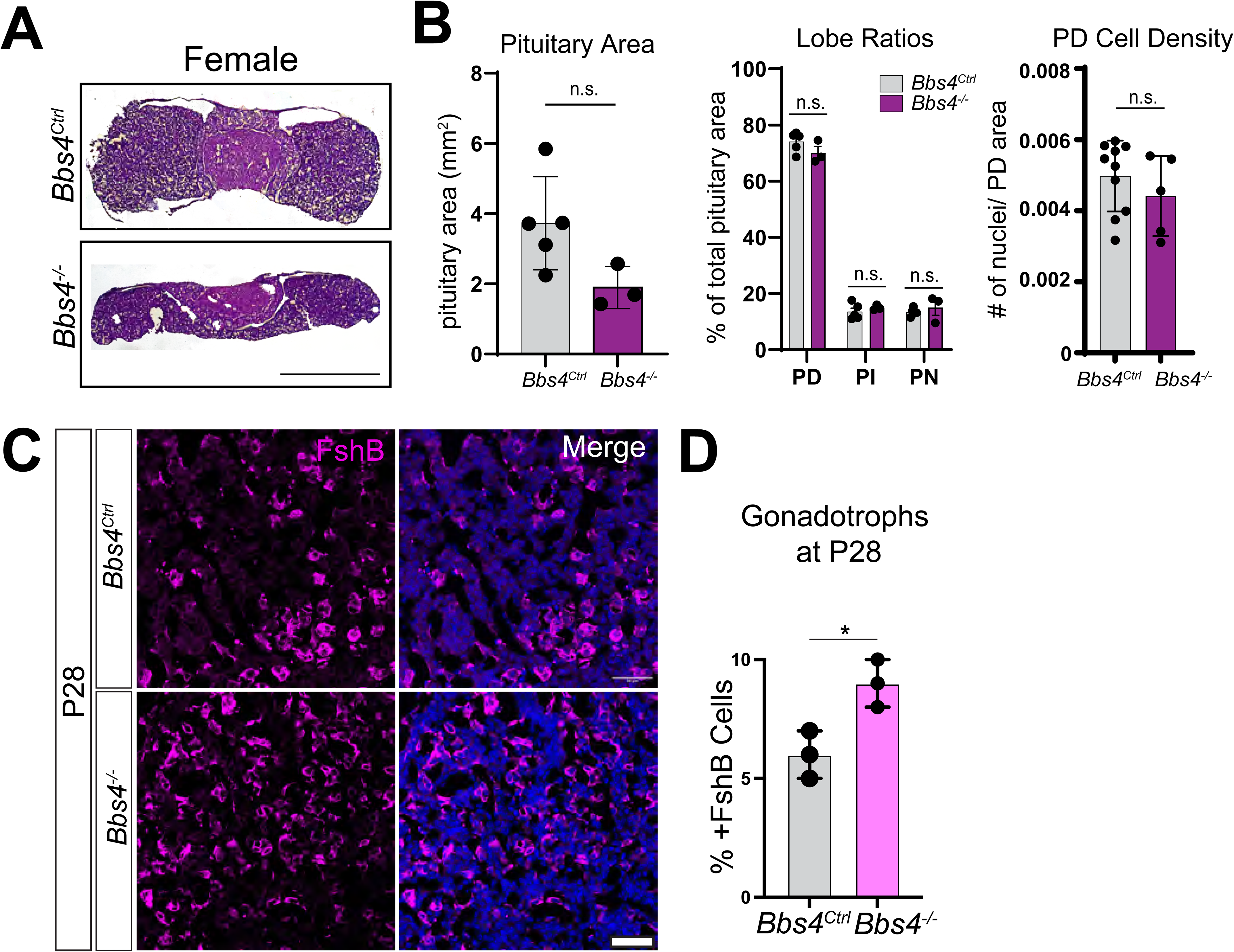
A. H&E staining of 8-week-old female pituitaries of mutants (*Bbs4^-/-^*) and controls (*Bbs4^Ctrl^*). Scale Bar 1 mm. **B.** Area and morphology analysis of *Bbs4^-/-^* mice compared to *Bbs4^Ctrl^* control females (student’s t-test). There is no significant difference in the area (student’s t-test), ratio of the 3 lobes of the pituitary (PD, PI, and PN) (one-way ANOVA) or PD cell density (student’s t-test). **C.** Immunofluorescence staining of hormones in the PD of P28 male controls and mutants for gonadotrophs (Fshβ, magenta) and hoechst nuclei blue. Scale bar 50 µm. **D.** Quantification of the percent gonadotrophs shows a significant increase by P28 in male mutants (student’s t-test).

**Supplemental Figure 2.**
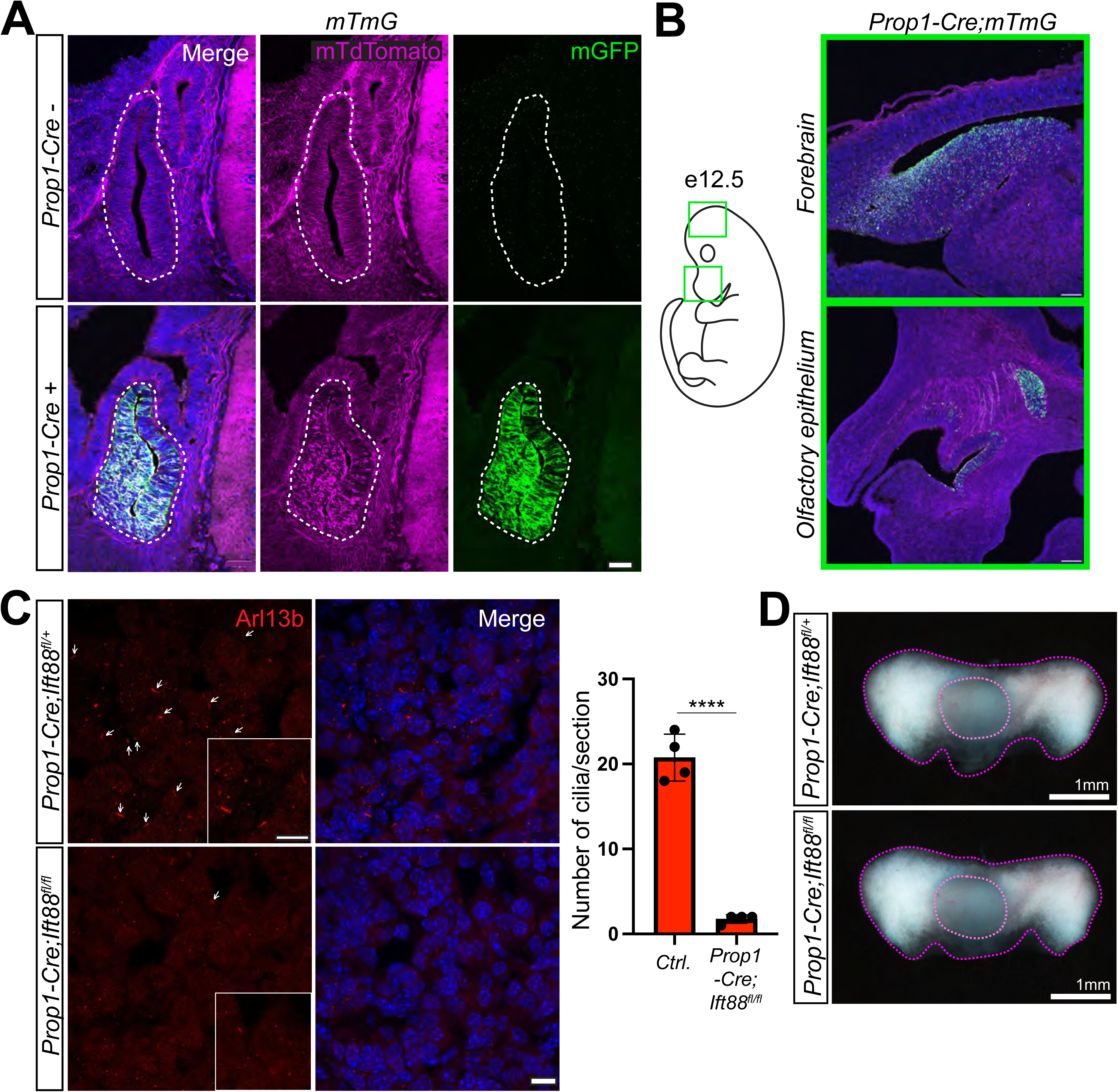
A. Immunofluorescence images of recombination of *mTmG* (membrane-tdTomato membrane-GFP) in *Prop1-Cre* positive and negative embryos at embryonic day 12.5. Green = GFP, Magenta = tdTomato, Blue = Hoechst. Scale bars 50 µm. **B.** Recombination present in 1 out of every 4 embryos in *Prop1-Cre mTmG* animals outside of the developing RP. Green = GFP, Magenta = tdTomato, Blue = Hoechst. Scale Bar 200 µm. **C.** Immunofluorescence images of cilia and corresponding quantification in *Prop1-Cre;Ift88^fl/+^* and *Prop1-Cre;Ift88^fl/fl^* confirming loss of cilia in pituitaries of *Prop1-Cre;Ift88^fl/fl^* animals at one month of age. Red = Arl13B, Blue= Hoechst. Scale bars = 10 µm and inset scale bars = 5 µm. **** = P-value <0.001. **D.** Brightfield images of *Prop1-Cre;Ift88^fl/+^*and *Prop1-Cre;Ift88^fl/fl^* pituitaries at one month of age. Scale bar = 1mm.

